# Human eukaryotic initiation factor 4G directly binds the 40S ribosomal subunit to promote efficient translation

**DOI:** 10.1101/2023.09.29.560218

**Authors:** Nancy Villa, Christopher S. Fraser

## Abstract

Messenger RNA (mRNA) recruitment to the 40S ribosomal subunit is mediated by eukaryotic initiation factor 4F (eIF4F). This complex includes 3 subunits: eIF4E (m^7^G cap binding protein), eIF4A (DEAD box helicase), and eIF4G. Mammalian eIF4G is a scaffold that coordinates the activities of eIF4E and eIF4A and provides a bridge to connect the mRNA and 40S ribosomal subunit through its interaction with eIF3. While the roles of many eIF4G binding domains are relatively clear, the precise function of RNA binding by eIF4G remains to be elucidated. In this work, we used an eIF4G-dependent translation assay to reveal that the RNA binding domain (eIF4G-RBD; amino acids 682–720) stimulates translation. This stimulating activity is observed when eIF4G is independently tethered to an internal region of the mRNA, suggesting that the eIF4G-RBD promotes translation by a mechanism that is independent of the m^7^G cap and mRNA tethering. Using a kinetic helicase assay, we show that the eIF4G-RBD has a minimal effect on eIF4A helicase activity, demonstrating that the eIF4G-RBD is not required to coordinate eIF4F-dependent duplex unwinding. Unexpectedly, native gel electrophoresis and fluorescence polarization assays reveal a previously unidentified direct interaction between eIF4G and the 40S subunit. Using binding assays, our data show that this 40S subunit interaction is separate from the previously characterized interaction between eIF4G and eIF3. Thus, our work reveals how eIF4F can bind to the 40S subunit using eIF3-dependent and eIF3-independent binding domains to promote translation initiation.

## INTRODUCTION

Messenger RNAs (mRNA) are selected for translation through recognition of key features including the 5′ 7-methyl-guanosine (m^7^G) cap and the 3′ poly(A) tail. Selection is mediated by the cap binding complex, eukaryotic initiation factor (eIF) 4F, and is a highly regulated step of translation initiation (1-5). eIF4F is composed of the m^7^G cap binding protein named eIF4E, a DEAD-box helicase named eIF4A, and a scaffold protein named eIF4G (reviewed in (6)). The eIF4G protein binds and coordinates the activities of eIF4E, eIF4A, and poly(A) binding protein (PABP). In mammals, mRNA recruitment to the 40S subunit is thought to be promoted by the direct interaction between eIF4G and the multi-subunit eIF3, which makes critical contacts with the 40S subunit. Consistent with this model, a truncated eIF4G that does not include the eIF3 binding domain cannot initiate translation in an eIF4G-dependent assay (7). A number of contacts between eIF4G, eIF3, and the 40S subunit have been observed in a recent cryo-EM model of the 48S complex, but it should be noted that the region surrounding the interaction of the eIF4F complex with eIF3 is of modest resolution (8). Following recruitment, the 40S subunit will scan along the mRNA in a 5′ to 3′ direction until it recognizes the start codon (usually AUG). Following this, initiation factors are released from the 40S and the 80S ribosome is formed to begin the elongation phase of translation. As a scaffolding protein, human eIF4G1 consists of a single 175 kDa polypeptide that contains several binding domains to promote mRNA recognition and translation (9,10). Conceptually, eIF4G can be divided into functional thirds (Fig. 1A). The N-terminal third of the protein is delineated by cleavage sites for foot-and-mouth disease virus (FMDV) L protease (L^pro^) and poliovirus 2A protease (2A^pro^). This region contains binding sites for eIF4E (9,11) and PABP (12,13). The C-terminal third of eIF4G binds the eIF4E kinase MAPK-activated protein kinase 1 (MNK1) (14) and has a second binding site for eIF4A (9,15-17). The central third of eIF4G extends approximately from amino acids 682-1104 (eIF4G_682-1104_). It contains RNA (18-20), eIF4A (15,17) and eIF3 (9,17) binding regions, and has been identified as the “functional core” of the protein, as defined by its ability to promote mRNA recruitment to the 40S subunit (21,22).

**Figure 1.**
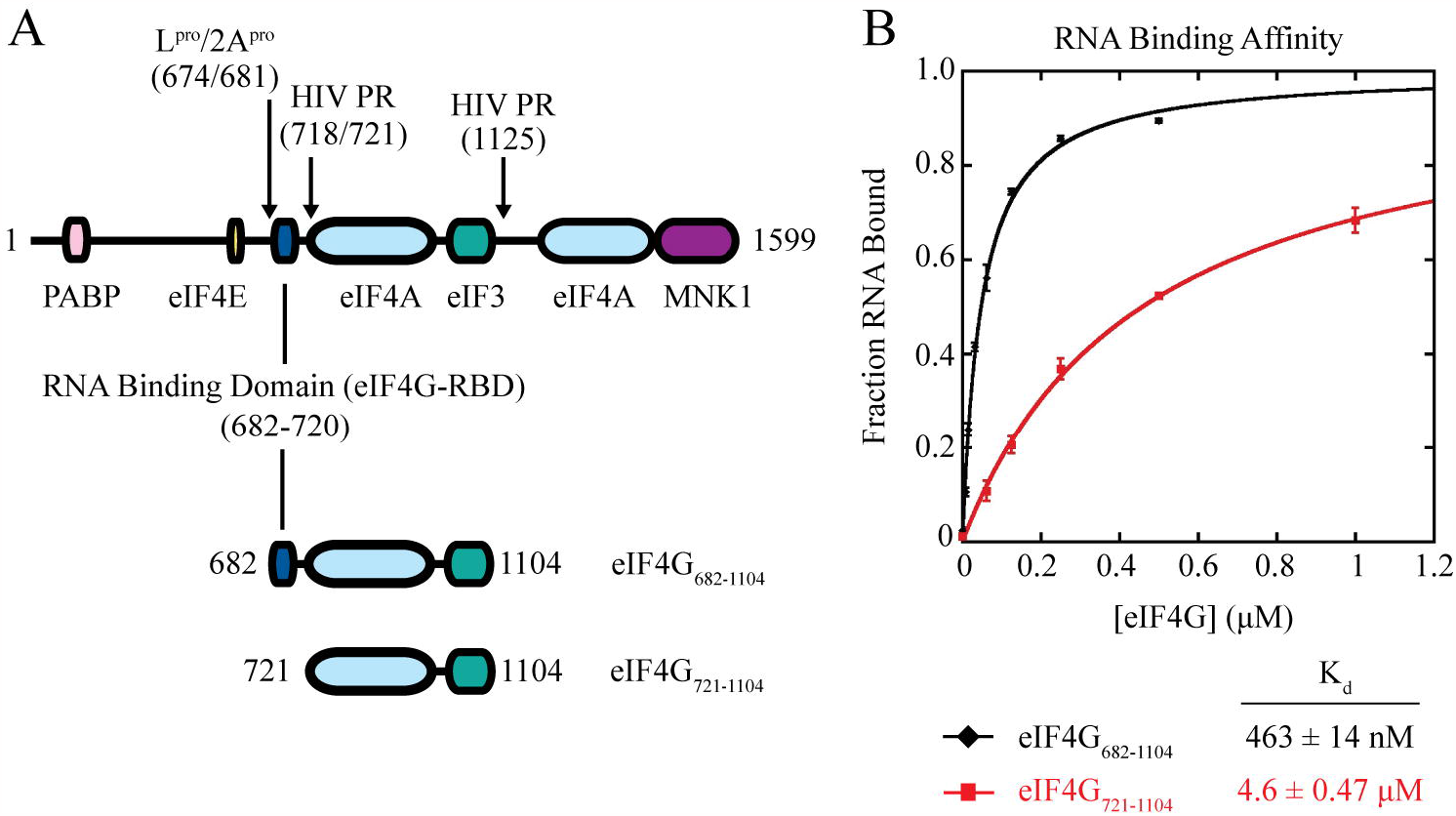
The eIF4G-RBD is required for high affinity RNA binding. (**A**), Domain map of full-length human eIF4GI and constructs used in this study. FMDV L protease (L^pro^), Poliovirus 2A protease (2A^pro^), and HIV-1 protease (HIV PR) cleavage sites are noted. The eIF4G-RBD (amino acids 682-720) is included in eIF4G_682-1104_ and truncated in eIF4G_721-1104_ to study the function of this domain. Each construct is tagged and mutated as noted in subsequent figures and legends. (**B**) Fluorescence polarization assays are used to determine the equilibrium dissociation constant (K_d_) of the eIF4G•RNA interaction using a 42 nucleotide 3′-end fluorescein-labeled RNA. The fraction of RNA bound by eIF4G_682-1104_ (black), or eIF4G_721-1104_ (red), is the average of at least 3 trials and error bars indicate the SEM.

The roles of the eIF4A and eIF3 binding regions in the eIF4G core have become established, but the precise role, and even the requirement for, RNA binding to this core domain remains unclear. Crosslinking between eIF4E and the mRNA cap show that eIF4G•eIF4E binding increases m^7^G cap association of eIF4E when recombinant eIF4G constructs added to the reaction contained the RNA binding region (23,24). Replacing the native RNA binding domain with that of the RNA recognition motif 1 (RRM1) of the La autoantigen showed that this effect was not sequence specific. Thus, the eIF4G interaction with mRNA may simply serve to tether eIF4E and increase its local concentration near the cap, thereby promoting m^7^G cap binding and translation (24). Nevertheless, one limitation of these experiments is that a crosslinking assay was used to monitor the formation of eIF4F on m^7^G capped mRNA. Thus, while these studies begin to explain the role of RNA binding as a tether to the mRNA, quantitative binding and translation assays are needed to fully characterize the function of this domain.

In *S. cerevisiae*, eIF4G contains 3 RNA binding domains, RNA1, -2, and -3, where RNA2 is in the same relative position (between the eIF4E and eIF4A binding sites) as the human eIF4G RNA binding domain. The eIF4G•eIF4E interaction was shown to not be essential in yeast, as eIF4G mutants impaired in this function were still viable. However, RNA2 deletion, eIF4E-binding double mutants were lethal despite retaining the ability to form eIF4G•PABP•mRNA complexes. These results suggest that RNA2 has an essential, non-redundant function beyond mRNA activation that is independent of the eIF4G•eIF4E interaction (25). While this RNA binding domain bears little sequence similarity between species, its comparable position within yeast eIF4G suggests that human eIF4G may bear similar, though yet undefined important functions. Throughout this manuscript, we will refer to this RNA binding domain in mammalian eIF4G1 as the eIF4G-RBD.

In mammalian cap-dependent translation, eIF4G bridges the mRNA and 40S subunit through its interactions with both eIF4E via its N-terminal binding domain and eIF3 via its binding domain in the eIF4G core. To promote viral protein synthesis during infection, both 2A^pro^ and L^pro^ cleave eIF4G to separate the N-terminal third from the middle and C-terminal domains, thereby reducing competition for the translation machinery from capped, endogenous mRNAs (26-28). Human immunodeficiency virus proteases (HIV PR) inhibit cellular translation by targeting eIF4G, most notably between the eIF4G-RBD and the eIF4A binding region (29,30). This results in a core domain containing only the eIF4A and eIF3 binding sites or a fragment that extends to the C-terminal end of the protein if the secondary site at amino acid 1125 is not cleaved (19,30). Protease cleavage of eIF4G in rabbit reticulocyte lysate (RRL) followed by centrifugation to pellet ribosomes and associated factors showed that L^pro^-cleaved eIF4G remains bound to ribosomes and eIF3 in the pellet, whereas eIF4G cleaved with HIV-2 PR, which removes the eIF4G-RBD, is released into the supernatant (19). These results indicate a possible role for the eIF4G-RBD of mammalian eIF4G in 48S complex formation, although it is also possible the effect was indirectly due to HIV-2 PR targeting other components of the RRL, such as eIF3 subunit d (eIF3d) which is known to bind eIF4G (7,19,31).

In this work, we investigate the specific role of the eIF4G-RBD. Using an eIF4G-dependent RRL system, we compare translation directed by 2 different eIF4G constructs: eIF4G_682-1104_, the eIF4G core that contains the eIF4G-RBD, and eIF4G_721-1104_, which is N-terminally truncated and does not contain the eIF4G-RBD (Fig. 1A). We find that while eIF4G_721-1104_ is capable of initiating translation, the eIF4G-RBD stimulates translation by about 1.5–2-fold. Unexpectedly, we identify a direct interaction between eIF4G and the 40S subunit. This interaction is dependent on the eIF4G-RBD, but independent of its role in promoting eIF4A-dependent helicase activity and tethering eIF4F to the mRNA. Quantitative binding assays we show that the eIF4G-RBD provides a direct binding site to the 40S subunit that is separate to the interaction between eIF4G and the eIF3 complex.

## RESULTS

### The eIF4G-RBD is required for high affinity RNA binding to eIF4G_682-1104_

Published studies have helped to locate and attribute general RNA binding properties to the human eIF4G-RBD (amino acids 682-720) (Fig. 1A) (18,20,24,32). The eIF4G-RBD has been proposed to help tether eIF4F to a cellular mRNA or a subset of factors to a viral transcript for ribosome recruitment. To determine the precise contribution that the eIF4G-RBD provides for RNA binding to the eIF4G core domain (eIF4G_682-1104_), we first used a steady-state fluorescence polarization assay to determine the equilibrium dissociation constant (K_d_) of a 42 nucleotide 3′-end fluorescein-labeled RNA (CAA-42-FL) and eIF4G in the presence (eIF4G_682-1104_) or absence (eIF4G_721-1104_) of the eIF4G-RBD (Fig. 1A). The change in fluorescence anisotropy that was specific to the eIF4G_682-1104_ or eIF4G_721-1104_ was measured, as described in *Experimental Procedures*. A strong anisotropy increase upon titration of either eIF4G construct was observed, indicating productive binding to form a complex (Fig. 1B). Converting anisotropy values into the fraction of CAA-42-FL bound at each eIF4G concentration yields a K_d_ of 463 ± 14 nM for eIF4G_682-1104_ and 4.60 ± 0.47 μM for eIF4G_721-1104_ (Fig. 1B; see Table 1 for *K*_d_ values). Thus, eIF4G_682-1104_ binds RNA with a ∼10-fold greater affinity than eIF4G_721-1104_, showing that the eIF4G-RBD provides a non-specific RNA binding affinity for the core eIF4G domain.

### The eIF4G-RBD promotes translation initiation independent of mRNA tethering

We next used an eIF4G-dependent translation assay in nuclease-treated RRL to determine the role of the eIF4G-RBD in translation. This assay was adapted from work by the Hentze lab, where an eIF4G “ribosome recruitment core” was identified *in vivo* to include amino acids 682-1131 that contains the eIF4G-RBD, the central eIF4A binding domain, and the eIF3 binding domain (21). Our lab showed that a similar eIF4G truncation (eIF4G_682-1104_) can efficiently recruit ribosomes to a transcript for translation using a RRL system (7). For this assay, a renilla luciferase reporter RNA with an engineered boxB hairpin in the 5′ UTR specifically recruits a 22 amino acid sequence of the bacteriophage λ transcription anti-terminator protein N (λN). Using the boxB hairpin, λ-eIF4G is directed to the reporter RNA independent of eIF4E, and ribosomes are recruited solely through interaction with λ-eIF4G. Luciferase translation is initiated by addition of recombinant purified λ-eIF4G to the lysate, and 5′-end dependent initiation is prohibited by the inhibitory hairpin placed at the 5′-end of the uncapped RNA (Fig. 2A). The boxB•λN interaction has a reported picomolar affinity, which ensures any λ-eIF4G proteins added to the RRL will be efficiently tethered at low concentrations to the RNA template to recruit ribosomes (7,21,33,34).

**Figure 2.**
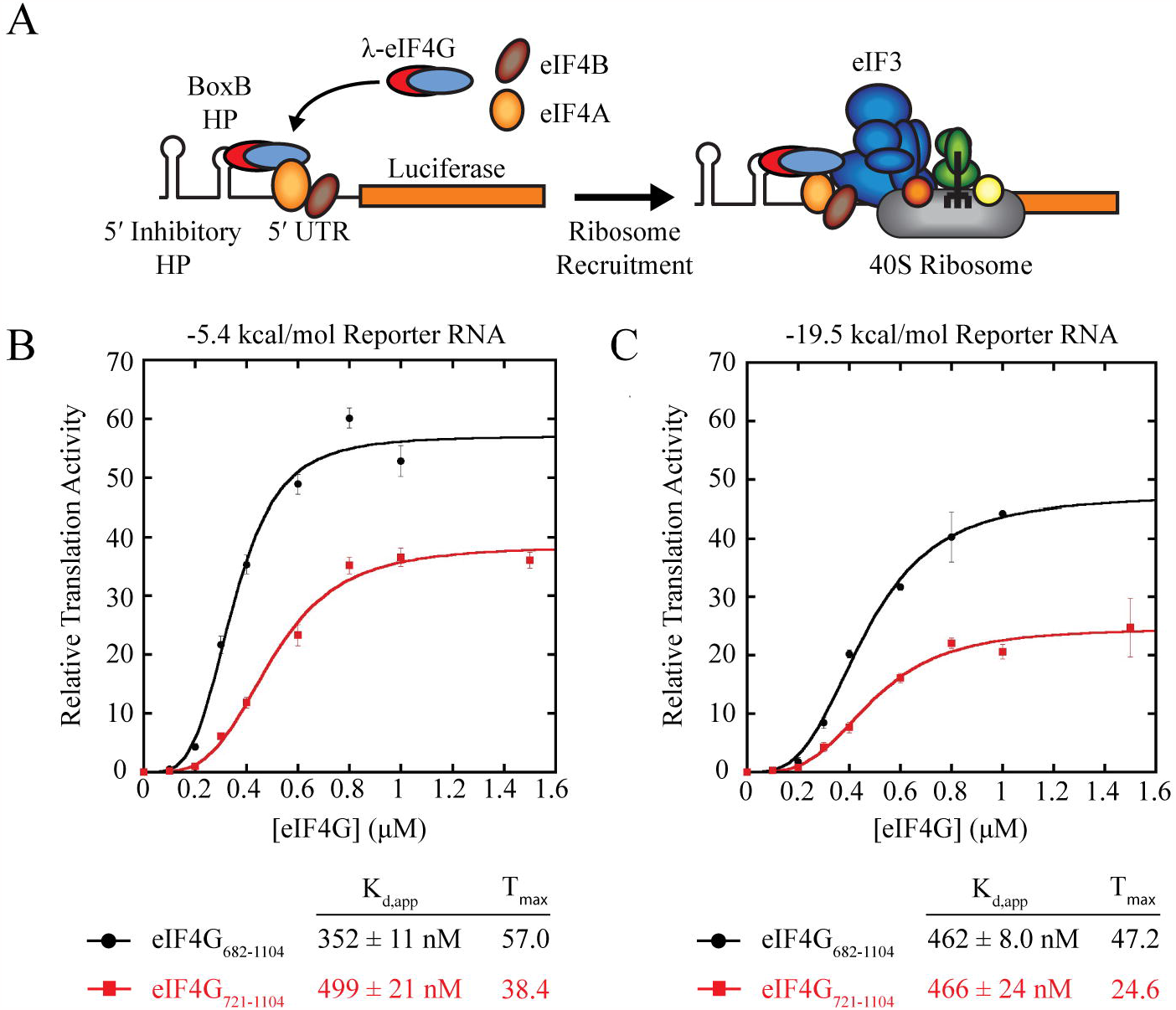
The eIF4G-RBD stimulates translation activity independent of mRNA tethering. (**A**) An eIF4G-dependent translation assay mediates reporter RNA recruitment to the ribosome independent of m^7^G cap binding. λN-tagged eIF4G constructs are recruited to a renilla luciferase reporter RNA via the boxB hairpin (BoxB HP) upstream of the 5′ UTR. An inhibitory stem loop near the 5′ end of the uncapped RNA prohibits 5′ end-dependent translation, and RNA recruitment to the ribosome occurs solely through interaction with λ-eIF4G. (**B**) and (**C**), Relative luciferase translation activity of 2 different RNA reporters with either low (**B**) or high (**C**) amounts of secondary structure in the 5′ UTR. λ-eIF4G_682-1104_ (black line) or λ-eIF4G_721-1104_ (red line) was titrated into nuclease-treated RRL and incubated for 30 min at 30 °C prior to measuring renilla luciferase activity, as described in *Experimental Procedures*. The K_d,app_ and maximum relative translation activity (“T_max_”) are the average of 3 trials and error bars indicate the SEM.

To test the independent role of the eIF4G-RBD in regulating translation, we compared translation activity of λ-eIF4G_682-1104_ versus the eIF4G-RBD truncation λ-eIF4G_721-1104_. We measured luciferase translation using increasing concentrations of λ-eIF4G to calculate the apparent equilibrium dissociation constant (K_d,app_) as well as the maximum relative translation activity (T_Max_) of each complex. Calculating K_d,app_ is needed to determine whether any differences in observed translation are due to differences in affinity or activity of each complex. Translation is normalized to a reaction containing no λ-eIF4G, and this background is subtracted prior to plotting relative translation as a function of λ-eIF4G concentration.

We tested our system using a reporter RNA with a relatively unstructured 5′ UTR (the region after the boxB hairpin), with a ΔG of -5.4 kcal/mol (Fig. 2B). Titration of λ-eIF4G_682-1104_ into the RRL increased the maximum relative translation activity by 57-fold over background (Fig. 2B). In contrast, titration of λ-eIF4G_721-1104_ increased the maximum relative translation activity by only 38.4-fold translation over background (Fig. 2B). Thus, the presence of the eIF4G-RBD enhances translation by ∼48 % compared to the eIF4G construct that does not possess the eIF4G-RBD. Fitting the increase in translation versus concentration of eIF4G added to a Hill equation (*Experimental Procedures*) reveals a K_d,app_ of 352 ± 11 nM for λ-eIF4G_682-1104_ and a slightly reduced K_d,app_ of 499 ± 21 nM for λ-eIF4G_721-1104_. While there is a modest reduction in the apparent affinity of translation complexes with the loss of the eIF4G-RBD, the reduction in maximum translation is clearly apparent. We note that translation activity was inhibited in the presence of >1 μM λ-eIF4G_682-1104_, while no such inhibition was observed using λ-eIF4G_721-1104_ (data not shown). Although the reason for this inhibition is not clear, it is possible that the eIF4G-RBD non-specifically inhibits translation when the RRL is supplemented with higher concentrations of λ-eIF4G_682-1104_.

We next tested if an RNA reporter containing a more structured 5′ UTR, with a ΔG of -19.5 kcal/mol, might have a different requirement for an eIF4G containing the eIF4G-RBD than the reporter with a 5′ UTR of -5.4 kcal/mol. Titration of λ-eIF4G_682-1104_ into the RRL increased the maximum relative translation activity by 47.2-fold over background (Fig. 2C). In contrast, titration of λ-eIF4G_721-1104_ increased the maximum relative translation activity by only 24.6-fold translation over background (Fig. 2C). Thus, the presence of the eIF4G-RBD enhances translation of this increased ΔG reporter by ∼92 % compared to the eIF4G construct that does not possess the eIF4G-RBD in this eIF4G-dependent assay. We calculated almost identical K_d,app_ of 462 ± 8.0 nM and 466 ± 24 nM for λ-eIF4G_682-1104_ and λ-eIF4G_721-1104_, respectively (Fig. 2C). These data therefore suggest that the eIF4G-RBD is not essential for mRNA translation in this eIF4G-dependent assay, but the amount of secondary structure located in the 5′ UTR does appear to modestly contribute to the degree of eIF4G-RBD-dependent translation that we observe; with a higher ΔG possessing a greater dependency on the eIF4G-RBD. Importantly, because the recombinant eIF4G proteins used are independently tethered to the mRNA via the λN tag, these experiments reveal a novel role of the eIF4G-RBD beyond simply acting as an mRNA tether.

### The eIF4G-RBD modestly increases affinity but not unwinding activity of the eIF4A helicase

Since we observe an effect of changes to the ΔG of the 5′ UTR on eIF4G-RBD-dependent translation, we explored the possible role of the eIF4G-RBD as an activator of eIF4A•eIF4B helicase activity using a fluorescence unwinding assay previously developed in our lab (35-37). For this assay, a 5′-Cy3 labeled RNA oligo is annealed to a template strand adjacent to a spectrally paired 3′-black hole quencher-labeled RNA. Strand separation by the helicase activity of purified eIF4A•eIF4B•eIF4G results in a dramatic increase in Cy3 fluorescence (Fig. 3A).

**Figure 3.**
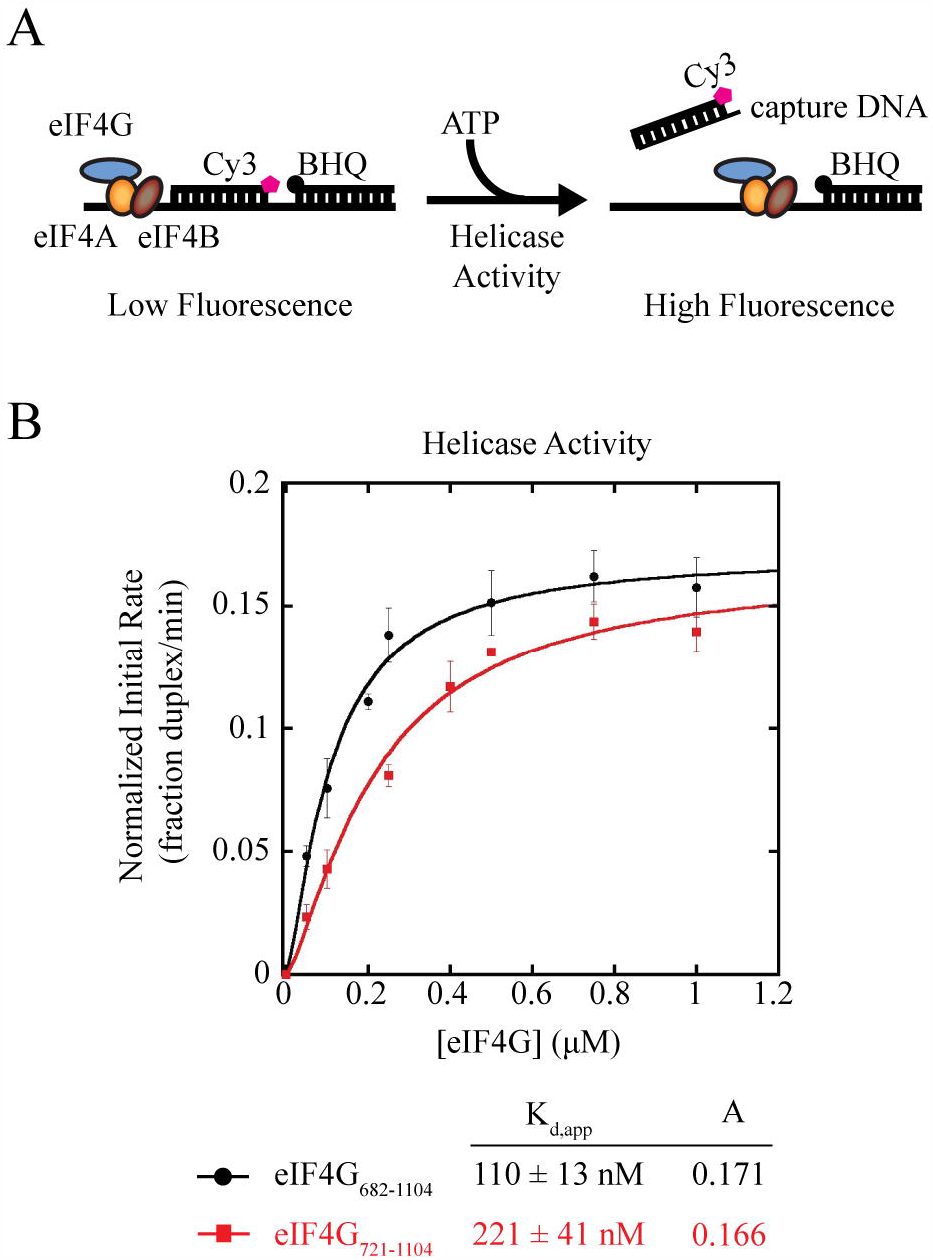
The eIF4G-RBD does not regulate the unwinding activity of the eIF4A helicase. **(A)** Schematic of eIF4A helicase fluorescent duplex unwinding assay. Briefly, a reporter RNA strand is modified on its 5′ end with Cy3 and annealed adjacent to a spectrally paired black hole quencher (BHQ) on a 3′-end modified RNA to a complimentary loading strand. Reporter RNA is incubated with 1 μM eIF4A, 1 μM eIF4B, and increasing amounts of eIF4G_682-1104_ or eIF4G_721-1104_. Addition of ATP/ MgCl_2_ starts the unwinding reaction, resulting in strand separation that is visualized as an increase in Cy3 fluorescence. A DNA capture strand prevents reannealing of the Cy3 reporter RNA. (**B**) The apparent equilibrium dissociation constant (K_d,app_) and maximum initial rates of unwinding (“A”) for the eIF4A•eIF4B•eIF4G complex containing eIF4G_682-1104_ (black line) or eIF4G_721-1104_ (red line) are calculated as previously described (see *Experimental Procedures*). Each data point represents the average of at least 3 experiments and error bars indicate the SEM, where initial rate of helicase activity (fraction of duplex unwound/min) is calculated from the linear portion of the unwinding curve for the eIF4G concentration indicated.

We used a saturating amount of eIF4A and eIF4B (1 μM as described in *Experimental Procedures*) and titrated in eIF4G_682-1104_ or eIF4G_721-1104_ to measure eIF4G-dependent duplex unwinding. The normalized initial rate of unwinding (fraction duplex unwound/min) was determined, as described in *Experimental Procedures*, and this was used to calculate the maximum rate of unwinding (A) and the K_d,app_ of the eIF4A•eIF4B•eIF4G•RNA unwinding complex (Fig. 3B). For eIF4G_682-1104_ we calculated a K_d,app_ of 110 ± 13 nM, while eIF4G_721-1104_ yielded a ∼2-fold greater K_d,app_ of 221 ± 41 nM (Fig. 3B). This modest difference in apparent affinity contrasts with the 10-fold affinity change of eIF4G•RNA in the absence of the eIF4G-RBD and reveals how these initiation factors cooperate with each other to form a stable unwinding complex (Fig. 1B). Interestingly, eIF4G_682-1104_ and eIF4G_721-1104_ had near identical maximum initial rates of unwinding, at 0.171 and 0.166 fraction duplex unwound/min, respectively (Fig. 3B). Thus, while there is a small deficiency in apparent affinity of the eIF4A•eIF4B•eIF4G_721-1104_•RNA complex, this can be overcome with higher concentrations of eIF4G_721-1104_ to yield a very similar maximum initial unwinding rate as eIF4G_682-1104_. In contrast, maximum relative translation activity was inhibited by ∼33-50% in the absence of the eIF4G-RBD (Fig. 2). Thus, these data suggest that the eIF4G-RBD is likely not promoting translation by coordinating the eIF4A helicase, at least not in the absence of the 40S subunit.

### eIF4G binds the 40S subunit independently of other initiation factors

We next inferred that if eIF4G is not promoting translation through coordination with the mRNA and helicase complex alone, it may be interacting directly or indirectly with the 40S subunit. To this end, we used a native gel electrophoresis assay to visualize eIF4G binding to the 40S subunit in the absence or presence of eIF3 and eIF4A.

We expressed and purified cysteine-free versions of eIF4G_682-1104_ and eIF4G_711-1104_, where all native cysteines have been converted to alanine, and where the residue E711 has been mutated to cysteine for site-specific modification with a Cy5 fluorophore to produce eIF4G-Cy5. The cysteine residues are not conserved in eIF4G, and the mutations do not affect eIF3 or eIF4A binding ((7) and data not shown). We note that 10 amino acids of the eIF4G-RBD, including 2 positively charged residues, remain in eIF4G_721-1104_ to allow fluorescent labeling at E711C. We confirmed both eIF4G-Cy5 constructs had similar levels of modification and were not degraded by the procedure using SDS PAGE (Fig 4A). We incubated 100 nM eIF4G-Cy5, 300 nM 40S subunits, 2 μM eIF4A, and 600 nM eIF3 in various combinations in the presence of 0.5 mM ATP-Mg on ice for 10 min followed by 30 °C for 5 min. Complexes were then separated on a 0.8% agarose gel to determine which factors were necessary for complex formation. Each gel was imaged for Cy5 visualization to track eIF4G and then stained with ethidium bromide to observe 40S subunits.

**Figure 4.**
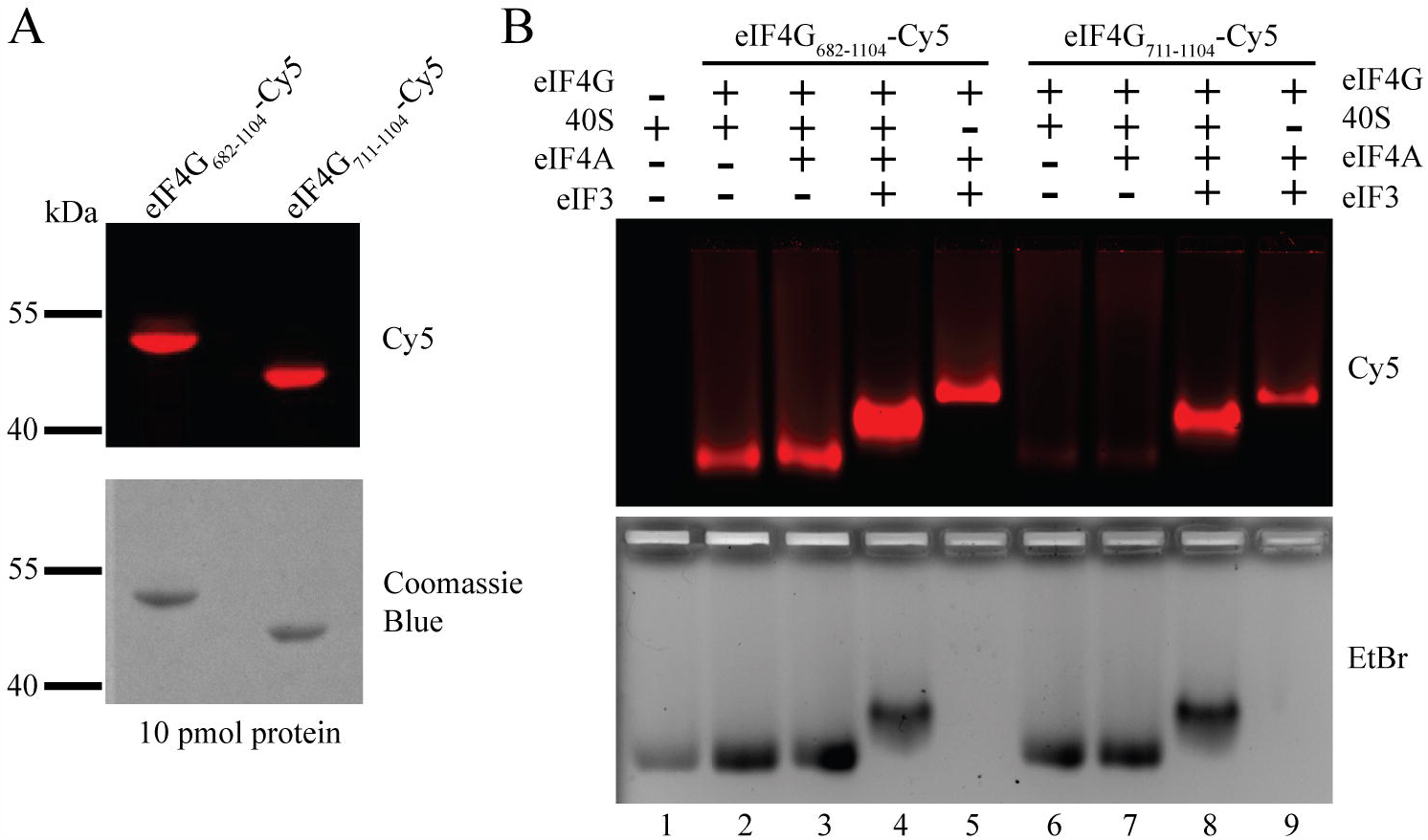
Native gel electrophoresis shows that the eIF4G-RBD stabilizes eIF4G binding to the 40S subunit in the absence of other initiation factors. **(A)** Cysteine-free eIF4G_682-1104_ and eIF4G_711-1104_ constructs each have a single cysteine mutation at residue E711 for site specific modification with Cy5. Following modification, 10 pmol of each protein was separated by SDS-PAGE, imaged to visualize the Cy5 fluorophore and stained with coomassie blue to confirm purity of each sample and equal levels of Cy5 modification efficiency. (**B**) 40S subunits (300 nM), eIF4A (2 μM), eIF3 (600 nM) and eIF4G-Cy5 (100 nM) were incubated together for 10 min on ice followed by 5 min at 30 °C in various combinations, then separated by native gel electrophoresis to visualize complex formation. Gels were imaged to visualize the eIF4G-Cy5 (upper panel) and stained with ethidium bromide to directly observe the 40S subunit (lower panel). The formation of eIF4G_682-1104_-dependent complexes (lanes 2-5) and eIF4G_711-1104_-dependent complexes (lanes 6-9) are shown. The outline of the wells are visible in the lower panel.

We found that eIF4G_682-1104_-Cy5 alone or with 2 μM eIF4A (a saturating amount based on previous studies (36,38)) largely remains in the well and does not enter the gel unless eIF3 is present (Fig. S1). Unexpectedly, eIF4G_682-1104_-Cy5 enters the gel and comigrates with the 40S subunit in the absence or presence of eIF4A (Fig. 4B). Because eIF4G_682-1104_ and eIF4A are relatively small compared to the 40S subunit, they do not substantially change the migration of the 40S subunit (Fig. 4B, lanes 1-3, top and bottom). Addition of eIF3 to form a 40S•eIF4A•eIF3•eIF4G_682-1104_-Cy5 complex appears to modestly enhance the interaction between eIF4G_682-1104_-Cy5 and the 40S subunit (Figure 4B, lane 4, top). As expected, the binding of eIF3 causes a sizeable upward shift of the 40S subunit (Fig. 4B, lane 4, bottom). The presence of apparent positive cooperativity between eIF4G_682-1104_-Cy5 and eIF3 in binding to the 40S subunit in this assay may reflect an increase in stability provided by the additional binding site that eIF3 possesses for eIF4G (7,39). In the absence of the 40S ribosomal subunit, we note that eIF4A•eIF3•eIF4G_682-1104_-Cy5 forms a stable complex (Fig. 4B, lane 5, top panel).

We then compared ribosome and initiation factor complex formation using eIF4G_711-1104_-Cy5, which lacks the majority of the eIF4G-RBD (10 amino acids remain to allow fluorescent modification). In this assay, we detect very minimal formation of a 40S•eIF4G_711-1104_-Cy5 or 40S•eIF4A•eIF4G_711-1104_-Cy5 complex (Figure 4B, lanes 6-7, top). The addition of eIF3 appreciably stabilizes eIF4G_711-1104_-Cy5 binding to the 40S subunit and resulted in a similar upward shift as seen previously (Fig. 4B, lanes 4 and 8). We confirmed eIF4G_711-1104_-Cy5 can still form a complex with eIF4A and eIF3 (Fig. 4B, lane 9), and that eIF4G_711-1104_ binding to eIF3 is unchanged in the presence of eIF4A (7). Taken together, we interpret the data to indicate that the eIF4G-RBD possesses an independent binding domain for the 40S subunit that is separate to the binding site on eIF4G for the eIF3 complex. As mentioned above, we note that there appears to be some degree of positive cooperativity between eIF4G and eIF3 binding to the 40S subunit, suggesting that the two binding sites may communicate with each other to result in maximum eIF4G binding affinity to the 40S subunit.

To extend these results and test if the second eIF4A binding domain in the C-terminal domain of eIF4G affects 40S subunit binding, we also tested eIF4G_682-1599_-Cy5 and eIF4G_711-1599_-Cy5 under the same conditions. We confirmed both eIF4G-Cy5 constructs had similar levels of modification and were not degraded by the procedure using SDS PAGE (Fig. S2A). The eIF4G_682-1599_-Cy5 construct directly binds the 40S ribosome in a similar way to eIF4G_682-1104_-Cy5 (Fig. S2B, lanes 2-5). Interestingly, we observe that the eIF4G-RBD truncation, eIF4G_711-1599_-Cy5, appears to bind the 40S subunit to an appreciably greater degree than eIF4G_711-1104_-Cy5 (Fig. S2B, lane 6, versus Figure 4B lane 6). While this binding is appreciably less than observed for eIF4G_682-1599_-Cy5, this weak binding may suggest an uncharacterized RNA binding region exists in the C-terminal third of eIF4G. Consistent with this, both eIF4G_682-1599_ and eIF4G_711-1599_ have a higher non-specific binding affinity for RNA in a fluorescence polarization assay than their C-terminal domain truncated counterparts (Fig. S2C and Fig. 1B). It is also possible that the C-terminal domain cooperates with the eIF4G core to promote its RNA binding affinity. Interestingly we note that upon addition of eIF4A, the 40S•eIF4G_711-1599_-Cy5 complex formation appears to be slightly reduced, only to be restored upon addition of eIF3 (Fig. S2B, lanes 7 and 8, top). This is a similar effect to that observed for eIF4A binding to eIF4G_711-1104_ (Fig. 4B, lanes 6-8). This small but reproducible anti-cooperative 40S binding by eIF4G and eIF4A likely reflects an altered conformation of eIF4G when bound to eIF4A. Taken together, these data show that eIF4G can stably and independently bind the 40S subunit through the eIF4G-RBD, and that this interaction is a separate binding site to the previously characterized eIF4G•eIF3 interaction.

### eIF3 enhances eIF4G recruitment to the 40S subunit independent of the eIF4G-RBD

One disadvantage of the native gel assay is that it is not an equilibrium binding assay. To gain a more rigorous understanding of the interaction between eIF4G and the 40S subunit, we quantitatively analyzed eIF4G binding to the 40S subunit using an equilibrium fluorescence polarization assay. To this end, we labeled the same single cysteine residue, E711C, with fluorescein to create eIF4G_682-1104_-Fl and eIF4G_711-1104_-Fl and used these to monitor binding to the 40S subunit. For these experiments, we focused on using an eIF4G/4A/ATP complex since we reasoned that it is the most physiologically relevant complex to characterize. Incubating increasing amounts of the 40S subunit with 20 nM eIF4G_682-1104_-Fl or eIF4G_711-1104_-Fl in the presence of a saturating amount of eIF4A and 0.5 mM ATP-Mg results in a K_d_ of 129 ± 9 nM and 563 ± 72 nM for eIF4A•eIF4G_682-1104_-Fl and eIF4A•eIF4G_711-1104_-

Fl respectively (Fig. 5A and 5B, dotted lines). Thus, there is a ∼4.5-fold reduction in affinity in the absence of the eIF4G-RBD, which is consistent with the reduced binding of eI4G in the absence of the eIF4G-RBD on the native gel assay. To explore the possible cooperative binding between the eIF4G-RBD and eIF3, we next added 200 nM eIF3 to each reaction in addition to the saturating amount of eIF4A and measured 40S affinity. This concentration of eIF3 results in ∼50% eIF4G-Fl bound and was chosen to give a reasonable estimate of the K_d_ of the 40S•eIF4A•eIF3•eIF4G-Fl complex. Upon addition of this sub-saturating amount of eIF3, we measured a K_d_ of 76 ± 4 nM for eIF4G_682-1104_-Fl and 248 ± 64 nM for the RNA binding truncation eIF4G_711-1104_-Fl (Fig. 5A and 5B, solid lines). We also tested this reaction in the presence of 600 nM eIF3, with similar results observed (data not shown). These data reveal that both eIF4G_682-1104_-Fl and eIF4G_711-1104_-Fl bind to the 40S subunit with a modest ∼2-fold increase in affinity in the presence of eIF4A and eIF3. Thus, the binding site provided by eIF3 appears to stabilize the eIF4G•eIF4A•40S complex independently of the eIF4G-RBD. One limitation of the anisotropy assay is that changes in fluorescence polarization are dependent on a large difference in size of the labeled molecule and binding partner, or a large conformation change upon binding. As more binding partners (especially eIF3) are added to eIF4G-Fl, we note that the total change in fluorescence polarization is appreciably reduced in our assay (Table 1). In addition, the total change in anisotropy for eIF4G_711-1104_-Fl is appreciably smaller than for eIF4G_682-1104_-Fl, making it more challenging to use this assay to study the binding of this protein to the 40S subunit (Table 1). Nevertheless, the anisotropy data show that an eIF4G/4A/ATP complex binds directly to the 40S subunit with reasonably high affinity and a clear reduction in 40S binding affinity is observed in the absence of the eIF4G-RBD.

**Figure 5.**
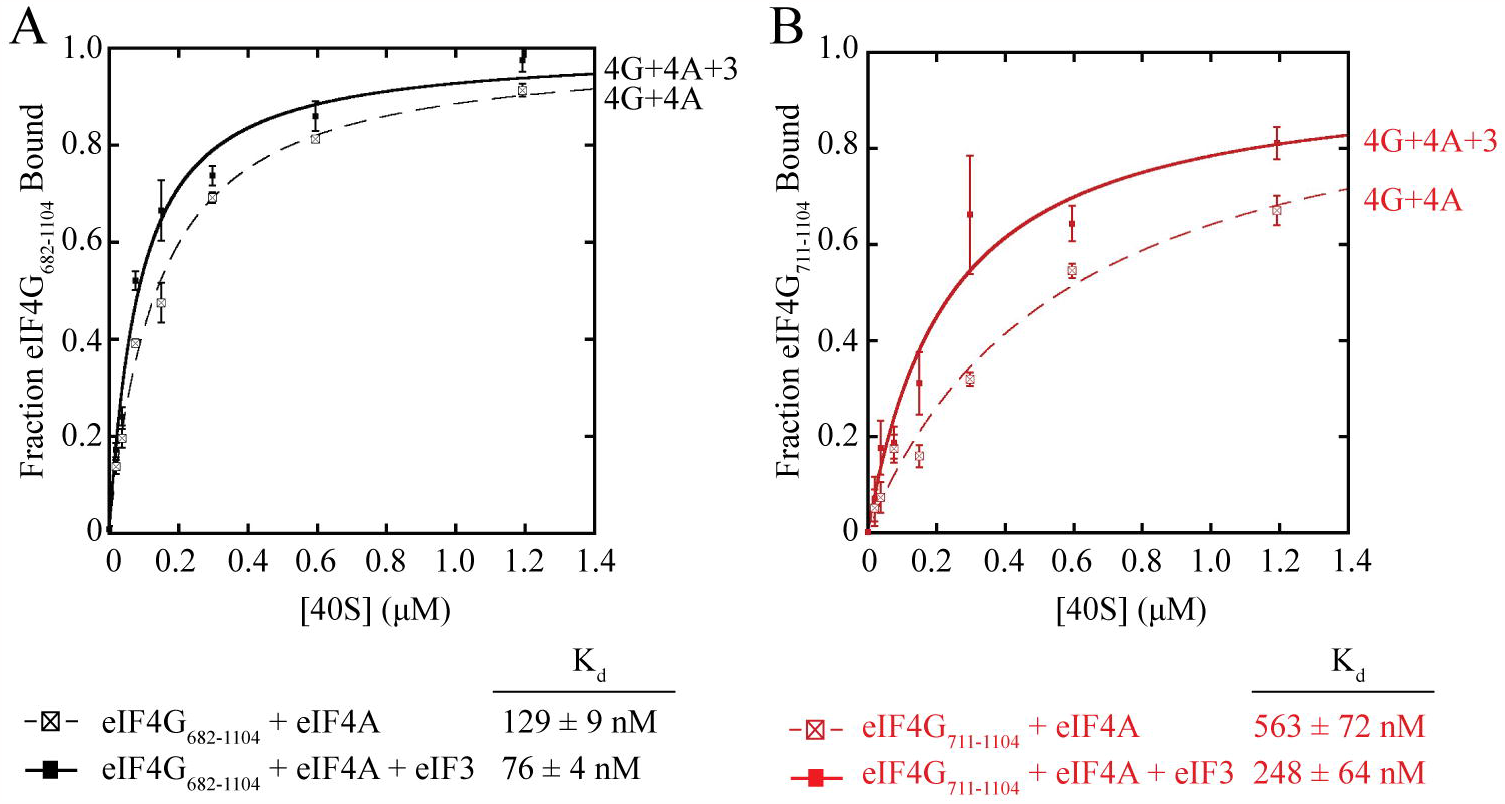
eIF3 enhances eIF4G recruitment to the 40S subunit independently of the eIF4G-RBD. **(A)** Equilibrium binding of eIF4G_682-1104_-Fl to the 40S subunit together with 3 μM eIF4A in the presence (solid black line) or absence (dashed black line) of 200 nM eIF3, as measured by the anisotropy assay. (**B**) Equilibrium binding of eIF4G_711-1104_-Fl to the 40S subunit together with 3 μM eIF4A in the presence (solid red line) or absence (dashed red line) of 200 nM eIF3. For each experiment, the fraction of eIF4G-Fl bound at different concentrations of 40S subunit is shown. The data are the average of at least 3 trials and error bars indicate the SEM. Equilibrium binding data were fit to determine the equilibrium dissociation constant (K_d_) of eIF4G binding to the 40S subunit, as detailed in the *Experimental Procedures*.

## DISCUSSION

In this work, we have characterized the human eIF4G-RBD, a lysine/arginine/proline-rich region shown previously to have general RNA binding capabilities and assumed to contribute to the eIF4F binding and recruitment of mRNA (18,20,24,32). To understand the function of RNA binding by the eIF4G-RBD, we used an eIF4G-dependent translation assay to compare translation efficiency of eIF4G constructs in the presence or absence of the eIF4G-RBD (Fig. 2). Our work shows that the eIF4G-RBD stimulates translation by up to 2-fold when eIF4G is directly tethered to an RNA transcript. Tethering eIF4G to the mRNA circumvents the need for the cap binding protein eIF4E and enables one to separate a possible role of eIF4G-dependent “tethering” of the mRNA from other potential RNA binding functions. As such, the function of the eIF4G-RBD appears to go beyond simple tethering of the mRNA to the eIF4F complex since the high affinity boxB•λN-tag interaction cannot compensate for the presence of the eIF4G-RBD in translation assays. Previous work showed that switching the eIF4G-RBD with the RRM of the La autoantigen (in the absence of the eIF4A and eIF3 binding domains) substituted for the eIF4G-RBD in promoting chemical crosslinking of eI4E with the m^7^G cap (24). While that experiment indicates that non-specific RNA binding is important for promoting eIF4E binding to the m^7^G cap, we note that those experiments were carried out using an eIF4G construct that did not include the middle domain or C-terminal domain. It is therefore unknown if switching the eIF4G-RBD for the RRM of the La autoantigen could promote translation in a similar way to the eIF4G-RBD.

Increasing the predicted ΔG of the 5′ UTR of the boxB reporter assay has a noticeably larger inhibitory effect on eIF4G-dependent translation in the absence of the eIF4G-RBD compared to its presence (Fig. 2). This raised the possibility that the eIF4G-RBD may function to coordinate the helicase activity of eIF4A. Using a kinetic duplex unwinding assay shows that the eIF4G-RBD does contribute slightly to the apparent affinity of the eIF4A•eIF4B•eIF4G unwinding complex, but it does not substantially increase helicase activity since both constructs possess a similar maximum initial rate of unwinding upon saturation (Fig. 3). These results therefore indicate an alternative role beyond mRNA tethering and eIF4A helicase activity for the eIF4G-RBD during translation initiation in mammals.

Next, we turned our attention from eIF4G binding mRNA to test for a potential interaction with the 40S subunit. Using native gel electrophoresis and fluorescence polarization revealed a novel direct interaction between eIF4G and the 40S subunit. Truncating the eIF4G-RBD appreciably reduced eIF4G/4A/ATP binding to the 40S subunit by over 4-fold, indicating that the eIF4G•40S interaction is strongly enhanced by the eIF4G-RBD even in the presence of eIF4A and ATP (Fig. 4B and S2). Interestingly, we recently showed that non-specific RNA binding by the eIF4F complex is regulated by the nucleotide-bound state of eIF4A (38). Here, we have only tested the contribution that eIF4A makes on eIF4G binding to the 40S subunit in the presence of ATP (Fig. 4 and 5). Future work will be needed to rigorously determine if the nucleotide-bound state of eIF4A regulates eIF4G binding to the 40S subunit using a kinetic assay similar to that used for studying the non-specific binding of eIF4F to RNA. We note that the current eIF4G-Fl anisotropy signal that we possess is rather low, so this would need to be optimized to enable such a rigorous kinetic analysis to be made.

The binding of eIF4G to the 40S subunit is enhanced by eIF3 in the presence or absence of the eIF4G-RBD (Fig. 4B, 5, and S2). This finding shows that eIF4G recruitment via eIF3 or the eIF4G-RBD are independent. While we note a modest degree of positive cooperativity is observed between these binding domains in the native gel assay, no such cooperativity was observed in the equilibrium binding assay. As mentioned above, a kinetic analysis of eIF4G binding to the 40S subunit and the relative contributions that the eIF4G-RBD and eIF3 binding domains have on this interaction will be a focus of a future study. We also note that in the absence of the eIF4G-RBD, eIF4G_711-1104_-Fl binds directly to the 40S subunit in the fluorescence polarization assay as opposed to the gel shift assay (Fig. 4 and 5). This may reflect the 10 amino acids (including 2 positively charged residues) of the eIF4G-RBD that remain in the constructs used for these assays to allow for fluorescent labeling. Additionally, because the fluorescence polarization assay is at equilibrium, this could maintain interactions that would otherwise dissociate during the native gel assay. Alternatively, this could reflect an additional interaction between the HEAT repeat domain of eIF4G and the 40S subunit that has not yet been observed in the high-resolution cryo-EM structure of the 48S complex (40).

Revealing a direct interaction between the eIF4G-RBD and 40S subunit is consistent with previous work from the Ohlmann lab showing that eIF4G in RRL co-sediments with a ribosome pellet in sucrose gradients when untreated or preincubated with L protease (19). Importantly, when RRL is preincubated with HIV-2 protease (HIV PR), the eIF4G-RBD domain is cleaved from the central eIF4A and eIF3 binding domains, resulting in the loss of eIF4G•40S co-sedimentation despite eIF3•40S co-sedimentation remaining unchanged (Fig 1A and (19)). This work revealed many interesting observations, but it was not able to reveal a direct interaction between eIF4G and the 40S subunit since the work was done in RRL where many other factors are present. Subsequent work showed that both eIF4G and its binding partner eIF3d can be targeted by HIV-1 PR, which in turn could have explained the loss of eIF4G•eIF3•40S co-sedimentation in RRL (7,31). Our recent cryo-EM reconstruction of the 48S complex indicates that the N-terminal domain of eIF3d is located close to the eIF4F complex (40). We therefore tested if eIF3d cleavage can reduce the eIF4G•eIF3 interaction using fluorescence polarization and crosslinking assays. Consistent with the 48S structural model, we show that eIF4G binds to the N-terminal 114 amino acids of eIF3d that remain bound to the eIF3 complex upon cleavage by HIV-1 PR, and no affinity change is observed between the cleaved and uncleaved eIF3 complex with eIF4G (Fig. S3). Overall, these data support the conclusion that loss of the eIF4G-RBD results in loss of 40S ribosomal subunit binding independent of the eIF4G•eIF3 interaction.

Our data shows that the eIF4G-RBD plays an important role in promoting efficient translation through direct binding with the 40S subunit. Thus, eIF4G can interact with the 40S subunit directly via its eIF4G-RBD and indirectly through the eIF3 binding domain (Fig. 6). This prompts the question of where on the 40S subunit the eIF4G-RBD binds. The cryoEM structure of the 48S complex during scanning places eIF4G near the mRNA exit channel, revealing that mRNA is recruited to the 40S ribosome through a “slotting” mechanism. Unfortunately, there is no indication of the location of the eIF4G-RBD in this structure. It is also not clear whether the eIF4G-RBD plays a role at a different stage of initiation, such as an early step in mRNA recruitment. The 48S complex structure does, however, place constraints on possible binding sites and future biochemical and structural studies may be able to precisely locate eIF4G-RBD-interacting residues or nucleotides on the 40S subunit (40).

**Figure 6.**
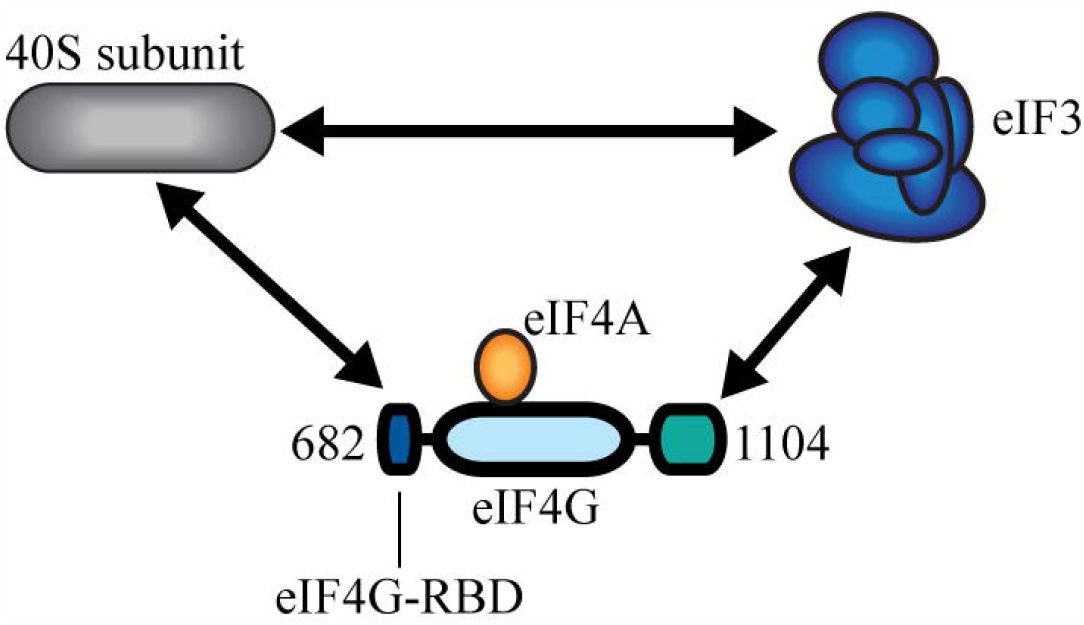
eIF4G binds to the 40S subunit through multiple contacts. Human eIF4G uses multiple interactions to form a molecular bridge that connects the eIF4F cap binding complex and the 40S subunit. An eIF3-specific binding domain (colored green) in human eIF4G indirectly stabilizes eIF4G on the 40S subunit. In this study, we characterize an unexpected direct interaction between the eIF4G-RBD (colored dark blue) and the 40S subunit that increases translation independently of mRNA tethering.

We previously characterized the presence of an autoinhibitory domain in human eIF4G that is relieved by the addition of eIF4E to promote eIF4A-dependent helicase activity (35,41). Interestingly, recent work has shown that the RNA2 domain in yeast eIF4G, which is positioned in a region analogous to the eIF4G-RBD, promotes the closed state of eIF4A, thereby increasing the RNA affinity and helicase activity of eIF4A. (42). While we do not observe any clear role of the eIF4G-RBD in promoting eIF4A helicase activity in our assays, it will be important for future work to determine if the 40S subunit binding function of the human eIF4G-RBD is conserved in RNA2 of yeast eIF4G. If RNA2 has a conserved 40S subunit binding function, it may help explain why an eIF3 binding domain in yeast eIF4G has not been observed.

The eIF4G scaffold is a multifunctional translation initiation factor. The role of the eIF4G-RBD in 40S subunit binding characterized in this work is in addition to the roles of eIF4G in eIF4F recruitment to mRNA and in duplex unwinding that is independent of the 48S complex. Detailed biochemical studies into mRNA recruitment and accommodation into the 40S subunit binding channel have revealed unexpected roles for eIF4G, eIF4A, and eIF4E in this process (43). Similar future studies may help determine whether the eIF4G-RBD plays a role in mRNA recruitment and/or scanning. Single-molecule studies comparing 48S complex formation and scanning with wild type or eIF4G-RBD mutant eIF4G proteins may also shed light on which steps of translation initiation are promoted by the eIF4G-RBD, as have been accomplished recently with the yeast translation initiation system (44). While our work has uncovered the importance of the eIF4G-RBD for translation, precisely how this novel interaction regulates mRNA recruitment, accommodation, or scanning, and whether it is targeted for translation regulation *in vivo* remains a topic for future work.

## EXPERIMENTAL PROCEDURES

### Recombinant expression of eIF4G constructs

– Recombinant constructs used in this work are denoted by the name of the factor, followed by the amino acid numbers it contains if it is a truncation of the wild type (WT) factor. All eIF4G constructs are for the human eIF4G1 protein. Numbering of eIF4G constructs is based on UniProt entry Q04637. Our recombinant eIF4G contains a glutamine insertion between residues 696 and 697, as shown in isoforms 7 and 8 (Q04637-7 and Q04637-8), but numbering will be consistent with isoform A as this currently referred to as the “canonical” sequence in other works.

All eIF4G constructs used in this work were expressed and purified from BL-21(DE3) cells, as previously described, including eIF4G_721-1104_ (incorrectly listed as eIF4G_711-1104_ in (7)). Purification of the eIF4G_682-1104_ Mut construct was accomplished using the same method as eIF4G_682-1104_. Purification λ-eIF4G and Bpa-labeled proteins was according to our previous study (7). Purification of eIF3, eIF4A, and eIF4B has been previously described (7,36).

### Fluorescence polarization assay

eIF4G•RNA affinity was measured using an uncapped 3′ fluorescent-labeled 42-nucleotide (42 nt) RNA as described previously (42-mer RNA-Fl) (38). Briefly, 20 nM 42-mer RNA-Fl was incubated with increasing amounts of eIF4G_682-1104_, eIF4G_721-1104,_ eIF4G_682-1599_, or eIF4G_721-1599_ (as indicated in figures) in Binding Buffer (20 mM Tris acetate (pH 7.5), 70 mM KCl, 2 mM free Mg2+ (supplemented as magnesium acetate), 0.1 mM spermidine, 1 mM DTT, 10% glycerol, and 0.1 mg/ml bovine serum albumin). Each binding reaction was allowed to reach equilibrium at 30 °C for 5 min prior to fluorescence polarization measurements in a Victor X5 Multilabel Plate Reader (Perkin Elmer). Each K_d_ measurement was calculated according to our previous work (45). Each K_d_ measurement is reported as the average of at least three experiments with error bars indicating the SEM.

eIF4G•40S ribosome affinity was measured using eIF4G_682-1104_, eIF4G_711-1104,_ eIF4G_682-1599_, or eIF4G_711-1599_, which had been expressed and purified with a single cysteine mutation at residue E711C and modified with fluorescein-5-maleimide (eIF4G-Fl). For each reaction, 20 nM eIF4G-Fl was incubated with an increasing amount of 40S subunit ± 3 μM eIF4A ± 200 nM eIF3 (as indicated in each figure) in Binding Buffer supplemented with 0.5 mM ATP/MgCl_2_. Each reaction was incubated on ice for 10 min, then 30 °C for 5 min and lastly 10 min at room temperature prior fluorescence polarization measurements. Each K_d_ measurement is reported as the average of at least three experiments ± standard error of the mean, and plots show the average of at least 3 experiments.

### BoxB tethered translation assay

The transcription template for the BoxB tethered assay is a derivative of our previously described template and was purchased from GenScript (35). The 5′ UTR was replaced with one of two 5′ UTR sequences possessing varying degrees of secondary structure:

- 5.4 Reporter RNA 5′ UTR (ΔG = -5.4 kcal/mol) 5′-CTCGAGACACCTACATTTGCTTCGAATATAACTGTGTCACTAGCAACCTCAAACAGA AGCTT-3′
- 19.5 Reporter RNA 5′ UTR (ΔG = -19.5 kcal/mol) 5′-CTCGAGACACCTACATTTGCTAGTGATATAACTGTGTCACTAGCAACCTCAAACAGA AGCTT-3′

The predicted stability of each 5′ UTR is determined by using the Predict a Secondary Structure Web Server by the D. H. Mathews lab at the University of Rochester Medical Center (46). Templates are amplified by PCR, *in vitro* transcribed, and purified using phenol-chloroform extraction following standard protocols. Reporter RNAs are diluted to 25 μM stocks in 20 mM Tris-acetate pH 7.5, 2 mM magnesium acetate, 100 mM KCl, and 0.2 mM DTT and folded by heating to 80°C, slow cooling to room temperature for ∼1.5 hours, then incubating for 10 min on ice (7).

Translation assays are carried according to our previous study (7). Briefly, 15 μl reactions contained 60% nuclease-treated rabbit reticulocyte lysate (Promega), 20 μM amino acid mixture minus leucine, 20 μM amino acid mixture minus methionine, 0.5 U/μl Recombinant RNasin Ribonuclease Inhibitor (Promega), 45 mM sodium chloride, 150 mM potassium acetate, 2 mM magnesium acetate, 0.25 μM uncapped RNA template, and λ-eIF4G at the concentrations specified in figures. These conditions minimize translation in the absence of λ-eIF4G. Reactions are incubated at 30 °C for 20 min prior to luminescence measurement for 10 s on the Victor X5 Multilabel Plate Reader (Perkin Elmer) by addition of 75 μl Renilla luciferase substrate (Promega). We normalized luciferase activity to a reaction containing no λ-eIF4G, and then subtracted this background before plotting the relative translation activity as a function of λ-eIF4G concentration. Plots in Fig. 2 are fit to the Hill equation:

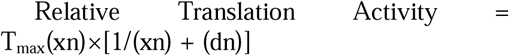

Where T_max_ is the maximum relative translation activity; d is the apparent K_d_, and n is the Hill coefficient. The average of 3 reactions ± standard error of the mean is presented here. All data are plotted using KaleidaGraph software (Synergy Software).

### Duplex unwinding assay

Unwinding reactions are carried out as previously described (35), using the same template. Briefly, a 24 nt cyanine 3 (Cy3)-labeled RNA reporter strand is annealed 1 nt upstream of a 19 nt black hole quencher (BHQ)-labeled RNA on an uncapped 64 nt RNA loading strand (Fig. 6A). These 3 RNAs are annealed at 500 nM in Unwinding Buffer (20 mM Tris-acetate pH 7.5, 2 mM magnesium acetate, 100 mM KCl, 0.2 mM DTT) by heating to 80 °C and slow cooling to room temperature for ∼1 hr. To prevent reannealing following helicase activity, a 24 nt DNA “capture” strand complementary to the Cy-3 reporter RNA is added at a 10X molar excess (2.5 μM) of the final RNA substrate concentration (250 nM). Unwinding reactions are carried out in 70 μl total volume containing 50 nM annealed substrate, 1 μM each eIF4A and eIF4B, eIF4G in the amount indicated in each figure, and 2 mM ATP/MgCl_2_ which is added after a stable baseline is reached to start the reaction. All reactions were carried out at room temperature, and data was fit to the Hill equation as described above and in previous works (35,36).

### Native gel electrophoresis

eIF4G•40S ribosome complex formation was visualized using eIF4G_682-1104_, eIF4G_711-1104,_ eIF4G_682-1599_, or eIF4G_711-1599_, which had been expressed and purified with a single cysteine mutation at residue E711C and modified with Cy5 (eIF4G-Cy5). For each reaction, 300 nM 40S subunits, 100 nM eIF4G-Cy5, 2 μM eIF4A and 600 nM eIF3 were combined as indicated in each figure in Binding Buffer supplemented with 0.5 mM ATP/MgCl_2_. Each reaction was incubated on ice for 10 min, then 30 °C for 5 min prior to loading on a 0.8% agarose gel in THEM Buffer (34 mM Tris acetate (pH 7.5), 57 mM Hepes (pH 7.5), 0.1 mM EDTA, 2 mM magnesium acetate) and separating for ∼5 hrs at 50 V. Gels were imaged to visualize the Cy5 fluorophore on eIF4G using an Azure Sapphire Biomolecular Imager (Azure Biosystems), then stained with ethidium bromide to directly observe the 40S subunit.

## Data availability

Data is presented within the manuscript and plasmids and other reagents are available for academic purposes upon request.

## Supporting information

This article contains supporting information.

## Acknowledgements

We wish to thank Masa Sokabe and Jessica Bolivar from the Fraser laboratory for many insightful comments and critical reading of the manuscript.

## Author Contributions

N.V. and C.S.F. conceived and designed the experiments and led the project. N.V carried out all experiments and both authors wrote and edited the manuscript.

## Funding and additional information

This work was supported by NIH grant R01 GM092927 to CSF. The content is solely the responsibility of the authors and does not necessarily represent the official views of the National Institutes of Health.

