## Supplemental Information for "Human eukaryotic initiation factor 4G directly binds the 40S ribosomal subunit to promote efficient translation"

**Supplementary Figure 1**

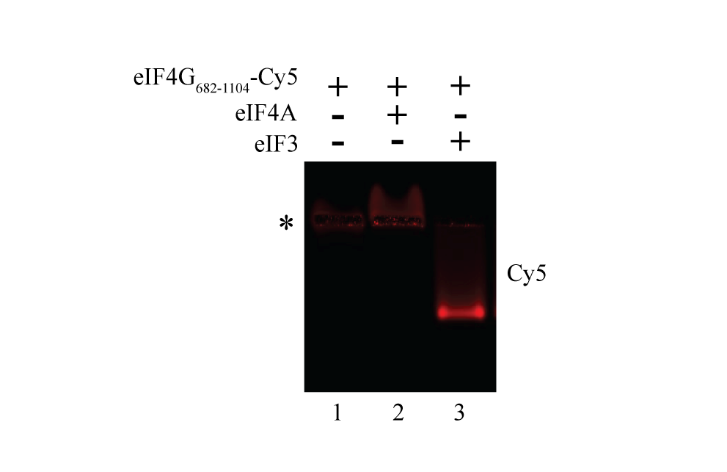

**Figure S1. Native gel electrophoresis controls for eIF4G_682-1104_-Cy5.** eIF4G_682-1104_-Cy5 largely remains in the well during native gel electrophoresis when free or bound to eIF4A (lanes 1 and 2). Upon addition of eIF3, eIF4G_682-1104_-Cy5 enters the gel and comigrates to form a distinct band indicating complex formation (lane 3). Location of the wells is marked by an asterisk (*).

Supporting Information:

Human eukaryotic initiation factor 4G directly binds the 40S ribosomal subunit to promote efficient translation

Nancy Villa and Christopher Fraser

**Supplementary Figure 2**

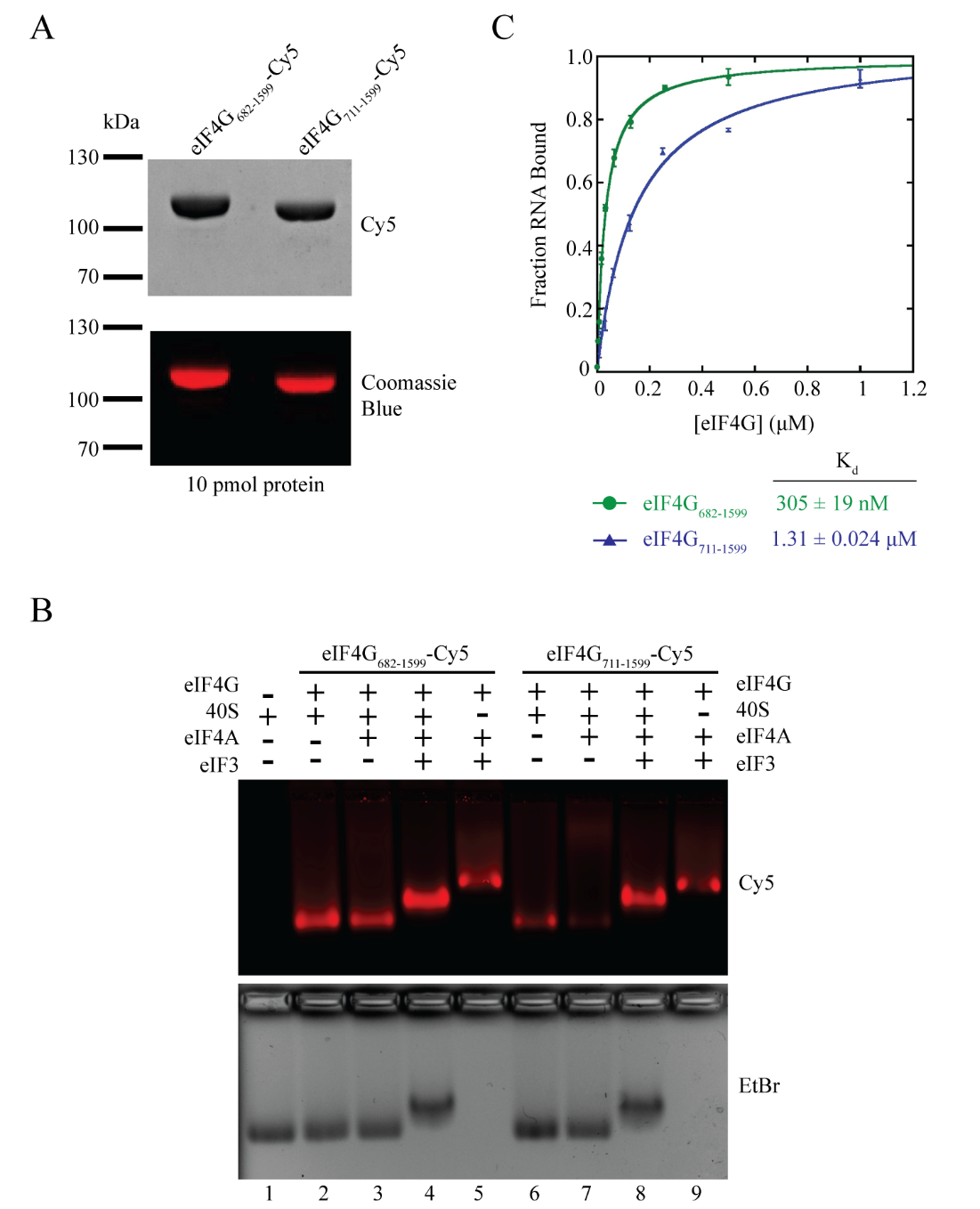

**Figure S2. eIF4G_682-1599_ binds the 40S subunit. (A)** Cysteine-free eIF4G_682-1599_ and eIF4G_711-1599_ constructs each have a single cysteine mutation at residue E711 for site specific modification with Cy5. Following modification, 10 pmol of each protein was separated by SDS PAGE, imaged to visualize the Cy5 fluorophore then stained with coomassie blue to confirm purity of each sample and equal levels of Cy5 modification efficiency. (**B**) 40S subunits (300 nM), eIF4A (2 μM), eIF3 (600 nM), 0.5 mM ATP, and eIF4G-Cy5 (100 nM) were incubated together for 10 min on ice followed by 5 min at 30 °C in various combinations, then separated by native gel electrophoresis to visualize complex formation. Gels were imaged to visualize the Cy5 fluorophore on eIF4G then stained with ethidium bromide to directly observe the 40S ribosomal subunit. Compare eIF4G_682-1599_ complex formation (lanes 2-5) to the RNA binding truncation eIF4G_711-1599_ (lanes 6-9). (**C**) Fluorescence polarization assays are used to determine the equilibrium dissociation constant (K_d_) of the eIF4G•RNA interaction using a 42 nucleotide 3′-end fluorescein-labeled RNA. Fraction of RNA bound by eIF4G_682-1599_ (green), or eIF4G_711-1599_ (blue). Data are the average of at least 3 trials and error bars represent the SEM.

Supporting Information:

Human eukaryotic initiation factor 4G directly binds the 40S ribosomal subunit to promote efficient translation

Nancy Villa and Christopher Fraser

**Supplementary Figure 3**

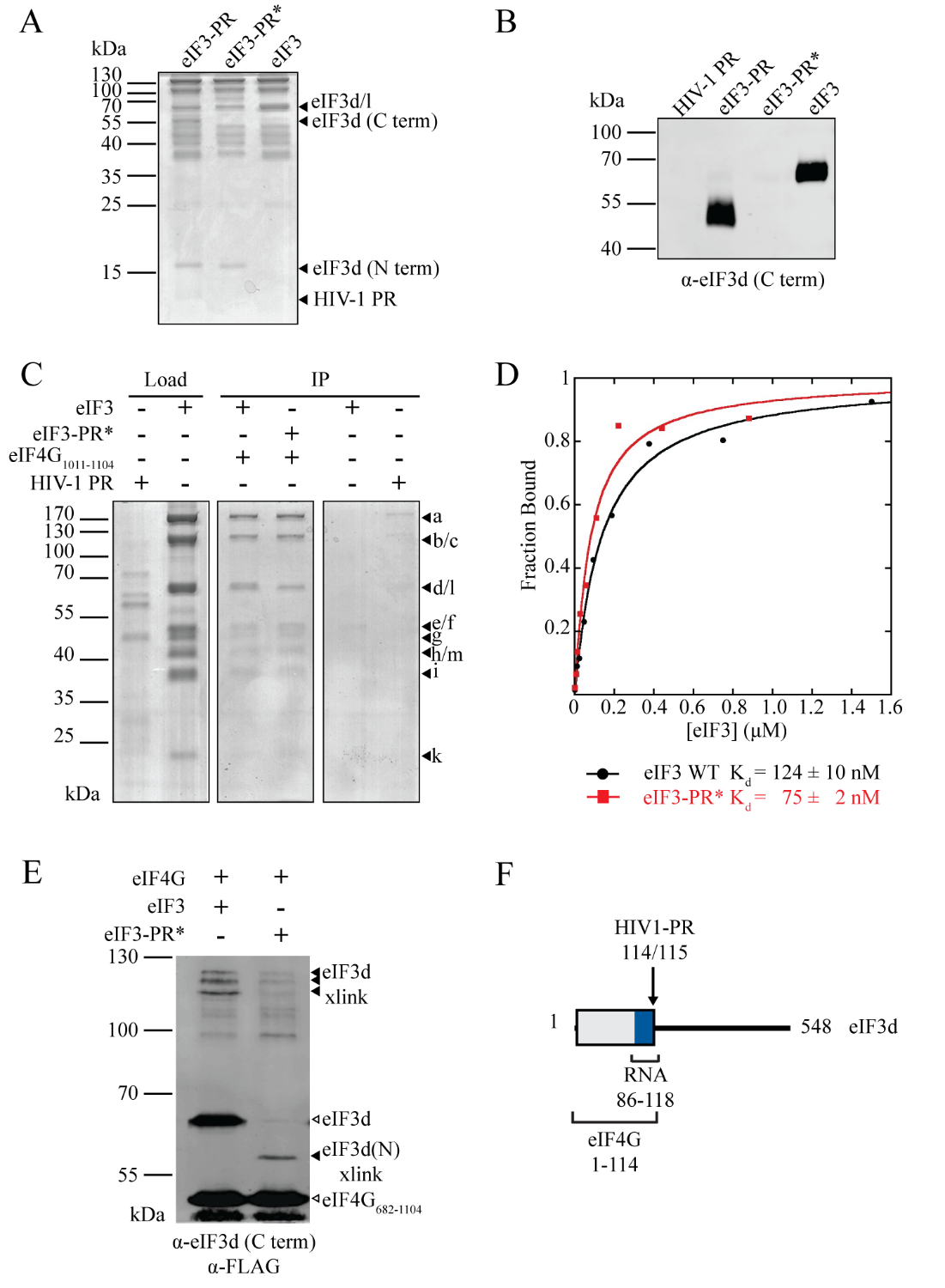

**Figure S3. Title. The N-terminal 114 amino acids of eIF3d binds eIF4G. (A**) Coomassie stained SDS-PAGE gel of eIF3d cleavage by HIV-1 Protease (HIV-1 PR). Asterisk (*) indicates that the sample was purified by gel filtration (as described in *supplemental experimental procedures*) following proteolysis, which results in dissociation of the eIF3d C-terminal domain from the eIF3 complex. (**B**) Western blot confirming eIF3d cleavage and removal of amino acids 115-548 from the eIF3 complex following size exclusion chromatography purification. (**C-D**) Fluorescence polarization and co-IP assays show eIF4G_1011-1104_, which contains the minimal eIF3 binding domain, binds both eIF3 and eIF3-PR* with approximately equal affinity. This indicates that eIF3d amino acids 1-114, which remain bound to the eIF3 complex, is sufficient for eIF4G binding. (**E**) Crosslinking with Bpa-labeled eIF4G_682-1104_ S1041X shows eIF4G binds to the N terminus of eIF3d following HIV-1 PR cleavage. (**F**) Model summarizing eIF3d subunit binding domains and cleavage sites.

Supporting Information:

Human eukaryotic initiation factor 4G directly binds the 40S ribosomal subunit to promote efficient translation

Nancy Villa and Christopher Fraser

**Supplementary Table 1**

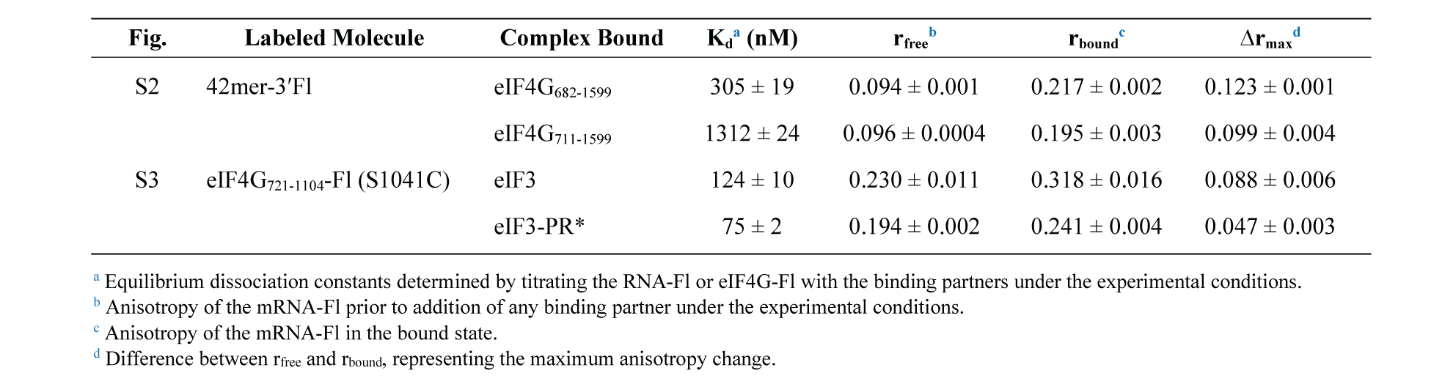

**Table S1. Summary of equilibrium binding parameters for Fig. S1.**

**SUPPLEMENTAL EXPERIMENTAL PROCEDURES**

*HIV-1 PR Expression and Proteolysis of eIF3*―A codon optimized construct of the HIV-1 protease (Strain NL4-3; HIV-1 PR) was purchased from GenScript. The sequence is shown below. HIV-1 PR was subcloned into a Pet28c vector with an N terminal 6X-His-MBP tag, expressed in BL-21(DE3) *E. coli* cells, and purified using a Ni-NTA column.

HIV-1 PR (Strain NL4-3)

MPQITLWQRPLVTIKIGGQLKEALLDTGADDTVLEEMNLPGRWKPKMIGGIGGFIKVRQYDQILIEICGHKAIGTVLVGPTPVNIIGRNLLTQIGCTLNF

Proteolysis reactions were carried out overnight at 4 °C with 3 mg/mL eIF3 and 2 mg/mL HIV-1 PR. The cleaved eIF3 complex was separated from HIV-1 PR by gel filtration through a Superose 6 (GE Healthcare) column in Gel Filtration Buffer (20 mM Hepes pH 7.5, 300 mM KCl, 10% glycerol, 1 mM DTT). The resultant eIF3 complex, eIF3-PR*, retained the N-terminal 114 amino acids of the eIF3d subunit while the C terminal portion and the eIF3j subunit dissociate at this salt concentration. This does not affect eIF4G•eIF3 complex binding affinity. We also discovered that the eIF3c subunit is cleaved by HIV-1 PR but only during gel filtration. This is likely an artifact of the *in vitro* reaction and purification and does not appear to influence eIF4G binding.

*eIF4G•eIF3 interaction assays*― eIF4G_1011-1104_ containing N-terminal 6X-HIS and C-terminal FLAG tags was used to coimmunoprecipitate eIF3 or eIF3-PR* using EZview Red Anti-FLAG M2 Affinity Gel (Sigma-Aldrich). Briefly, 10 μg eIF4G_1011-1104_ in 50 μl binding buffer (BB: 20 mM Hepes pH 7.5, 100 mM KCl, 10% glycerol, 0.5 mM DTT) were pre-incubated with the resin, and the excess was washed twice with 70 μl Wash Buffer (WB: 20 mM hepes pH 7.5, 200 mM KCl, 0.5% TritonX-100, 0.5 mM DTT, 5% glycerol). eIF3 (12 μg) was pre-incubated with HIV-1 PR or with binding buffer to cleave eIF3d, then added to the eIF4G-bound anti-FLAG resin. Once bound, excess eIF3 was washed away 3 times with 40 μl WB, and eluted in 20 μl of 250 ng/μl FLAG peptide. Results were analyzed by SDS PAGE.

For fluorescence polarization assays, all cysteine residues in eIF4G_721-1104_ were mutated to alanine by site directed PCR mutagenesis, and a single cysteine was introduced at S1041 for fluorescein labeling. Amino acids mutated were not conserved and did not appear to affect eIF4A or eIF3 binding functions of this protein. Labeling and fluorescence polarization assays were carried out as described previously (7).

The cotranslational benzophenone crosslinker incorporation and expression of eIF4G using amber mutants has been previously described in detail (7). Briefly, the unnatural amino acid p-Benzoylphenylalanine (Bpa; BaChem) is incorporated into eIF4G_682-1104_ cotranslationally in BL-21 (DE3) *E. coli* at S1041 using the pEVOL plasmid generously provided by Dr. Peter Schultz (The Scripps Institute). eIF4G_682-1104_-Bpa was purified and used in crosslinking reactions with intact eIF3 or eIF3-PR* complexes as described previously (7).
