## Supplementary material for "Human eukaryotic initiation factor 4G directly binds the 40S ribosomal subunit to promote efficient translation": Table 1

| **Fig.** | **Labeled Molecule** | **Complex Bound** | **K_d_^a^ (nM)** | **r_free_^b^** | **r_bound_^c^** | **∆r_max_^d^** |
| --- | --- | --- | --- | --- | --- | --- |
| 1 | 42mer-3′Fl | eIF4G_682-1104_ | 463 ± 14 | 0.097 ± 0.003 | 0.240 ± 0.001 | 0.143 ± 0.003 |
|  |  | eIF4G_711-1104_ | 4605 ± 467 | 0.100 ± 0.001 | 0.232 ± 0.006 | 0.132 ± 0.005 |
| 5 | eIF4G_682-1104_-Fl | eIF4A + 40S | 129 ± 9 | 0.133 ± 0.002 | 0.176 ± 0.001 | 0.043 ± 0.004 |
|  |  | eIF4A + eIF3^e^ + 40S | 76 ± 4 | 0.152 ± 0.002 | 0.182 ± 0.002 | 0.030 ± 0.002 |
|  | eIF4G_711-1104_-Fl | eIF4A + 40S | 563 ± 72 | 0.129 ± 0.001 | 0.156 ± 0.002 | 0.027 ± 0.001 |
|  |  | eIF4A + eIF3^e^ + 40S | 248 ± 64 | 0.144 ± 0.001 | 0.155 ± 0.001 | 0.011 ± 0.002 |

^a^ Equilibrium dissociation constants determined by titrating the RNA-Fl or eIF4G-Fl with the binding partners under the experimental conditions.

^b^ Anisotropy of the mRNA-Fl prior to addition of any binding partner under the experimental conditions.
^c^ Anisotropy of the mRNA-Fl in the bound state.

^d^ Difference between r_free_ and r_bound_, representing the maximum anisotropy change.

^e^ A subsaturating concentration, 200 nM, of eIF3 was used to estimate K_d_ changes in the presence of eIF3.

**TABLE 1. Summary of equilibrium binding parameters.**
